# Antimitotic chemotherapy promotes tumor NF-kB secretory phenotype and immunosuppressive CXCR2+ neutrophils chemotaxis in triple-negative breast cancers

**DOI:** 10.1101/2024.12.21.629876

**Authors:** Florian Chocteau, Fabien Gautier, Vanessa Josso, Elise Douillard, Philippe Juin, Sophie Barillé-Nion

## Abstract

Integrated approaches that help understand how tumors, as immune-surveilled ecosystems, respond to chemotherapy are crucial for developing effective antitumor treatments. We previously showed, in immunodeficient context, that antimitotic chemotherapy induced cGAS/STING pathway amplifying antitumor response through a paracrine IFN-1 secretome. We herein studied tumor progression and response to treatment using an immunocompetent murine model. scRNAseq analysis revealed that paclitaxel treatment altered tumor cell phenotypes, favoring tumor cells with a gene expression signature indicative of active NF-κB pathway with secretory phenotype. Treatment coincidently reduced IFN-I signature cells during tumor progression. The resulting secretory shift correlated with neutrophil recruitment to the tumor, particularly CXCR2+ neutrophils, thereby contributing to an immunosuppressive microenvironment. Pharmacological inhibition of CXCR2 receptor by navarixin reactivated antitumor immunity, enhancing NK cell infiltration and tumor cytotoxicity. Navarixin combination with paclitaxel significantly reduced tumor volume and metastasis. Targeting the NF-κB-driven secretory phenotype, in particular through neutrophil modulation, holds promise for improving TNBC treatment outcomes.

## INTRODUCTION

Triple Negative Breast Cancer (TNBC) remains a significant clinical challenge due to its aggressive nature and limited targeted treatment options (Sandarenu et al., 2022; Balic et al. 2023). Among the therapeutic strategies, antimitotic chemotherapy stands out as a cornerstone in managing TNBC but most patients experience tumor cell resistance with disease progression (Yin et al., 2020).

Resistance to antimitotic chemotherapy in TNBC is intricately linked to both intrinsic tumor resistance mechanisms and direct or tumor-dependent impact on the tumor microenvironment, particularly immune cells. TNBC tumors often exhibit intrinsic resistance to chemotherapy due to a variety of factors, including mutations in key genes involved in DNA damage repair pathways, activation of pro-survival signaling pathways (Nedeljković et al., 2019; Zheng et al. 2023). Moreover, chemotherapy-induced immunomodulation can further exacerbate resistance by promoting an immunosuppressive tumor microenvironment characterized by the recruitment of regulatory T cells (Tregs, Dabrowska et al., 2023) or myeloid immunosuppressive cells (Haist et al., 2021, Kersten et al., 2022), the release of Neutrophil Extracellular Traps (Shahzad et al., 2022), as well as the upregulation of immune checkpoint molecules such as programmed death-ligand 1 (PD-L1, Filderman et al., 2024). While the latter can be pharmacologically targeted through immune checkpoint inhibition, which has shown significant success in the clinic, the former remains a major obstacle. Consequently, the development of therapeutic strategies to overcome resistance in TNBC requires a multifaceted approach that targets both intrinsic tumor resistance mechanisms and modulates the immune microenvironment to enhance anti-tumor immune responses

Central to the immune response against tumors under chemotherapeutic pressure is the cGAS/STING pathway, a key surveillance mechanism for detecting cytosolic DNA. Upon activation by cytoplasmic DNA, cGAS generates cyclic GMP-AMP (cGAMP), which in turn activates STING, leading to the production of type I interferons (IFN-I) and NF-kB proinflammatory cytokines, playing a crucial role between tumor cells and innate and adaptive immunity (Guo et al., 2024; Jeltema et al. 2023; Yum et al., 2021). However, recent studies have uncovered a dichotomous role of STING in TNBC, playing both antitumor and protumor effects. In addition to promote spreading of chemo-induced tumor cell death (Lohard et al., 2020), STING activation enhances the priming and activation of dendritic cells, cytotoxic T cells or NK cells, promoting tumor cell elimination (Gan et al., 2022; Holicek et al., 2024). Conversely, chronic STING activation favors NF-κB pathway, associated with immune suppression and tumor progression, highlighting its complex role in tumor immunity (Hong et al., 2022; Li et al., 2023). Indeed, this NF-κB signaling pathway, favors the extracellular release of key factors in tumor progression, including chemokines, growth factors, and enzymes, which prevent effector cells of antitumor immunity from infiltrating the tumor and promote tumor invasion and migration (Hänggi et al., 2024; Lanng et al., 2024).

Thus, the interplay between antimitotic chemotherapy, the cGAS/STING and NF-κB pathways represents a dynamic and multifaceted aspect of TNBC treatment. While chemotherapy-induced immunogenic cell death can stimulate anti-tumor immune responses via STING activation, the context-dependent nature of STING signaling underscores the need for precise modulation in therapeutic strategies. Morevover, it is still unclear whether signaling pathways other than STING, notably NF-κB, might modulate immune response to treatment. Elucidating the nuances of different secretomes downstream of interconnected yet independent transcriptional pathways offers promise for improving the efficacy of TNBC treatment and harnessing the potential of immunotherapy to combat this aggressive disease. In this study, we examined the mechanisms underlying chemotherapy-induced secretory phenotypes downstream of STING and NF-κB and their influence on the tumor microenvironment in an immunocompetent context at all stages of TNBC progression.

## MATERIALS AND METHODS

### Cell lines

The 4T1-Luc2 cell line was obtained from ATCC and cultured in Dulbecco’s Modified Eagle Medium (DMEM) from Gibco (Saint Aubin, France), supplemented with 5% Fetal Bovine Serum (FBS) sourced from Eurobio (Courtaboeuf, France), 2 mM glutamine, and 1% penicillin/streptomycin from Gibco. CRISPR-Cas9-induced knockout (KO) cell lines targeting mouse genes were generated using specific single guide RNA sequences designed with the CRISPR design tool (http://crispor.tefor.net). The guide sequences (CACCTAGCCTCGCACGAACT for *Sting1* gene and CAGCGCAGCTCATTGGCGAT for *Cxcl1* gene) were inserted into the plentiCRISPRV2 vector (Addgene plasmid # 52961). Cells were selected using either 10 μg/ml puromycin (CRISPR^STING^) or 100 μg/ml hygromycin (CRISPR^CXCL1^), and knock-outs were confirmed through immunoblotting or ELISA analysis. *In vitro* treatments were administered at the following concentrations: 100 nM Paclitaxel from Sigma-Aldrich (St Quentin Fallavier, France), 0.2 nM Navarixin from MedChemExpress (Monmouth Junction, USA), and 10 μM AS602868 (Frelin et al., 2003).

### Biochemical assays

For immunoblot analysis, protein lysates were obtained from cells with RIPA buffer (89901, Thermofisher scientific) before separation on SDS-PAGE and transfer on nitrocellulose membranes. Membranes were then incubated for 1h at room temperature with the following primary antibodies (1/1000): Actin (MAB1501) from Millipore; Firefly Luciferase (ab21176), BCL2 (ab117115), BCL-xL (ab32370) from Abcam; HSP90 (610418) from BD Biosciences; IkBα (5483), pSer32-IkBα (9242), MCL-1 (94296), STING (13647) from Cell Signaling. Then membranes were incubated with the appropriate secondary antibodies for 1h at room temperature. Immobilon Forte Western HRP substrate kit (WBLUF0100, Millipore, Marne la Coquette, France) was used for immunoblot revelation on the Fusion FX + system (Veber).

For cytokine array, levels of different cytokines in cell supernatants were semi-quantitatively determined according to the manufacturer’s protocol (Mouse Cytokine Array C3, RayBiotech) on the Fusion FX + system (Veber).

For ELISA assays, levels of different chemokines in cell supernatants were determined according to the manufacturer’s protocols using following kits: CXCL1 (Mouse CXCL1 Sandwich ELISA Kit, Proteintech) and CCL5 (Mouse CCL5 Sandwich ELISA Kit, Proteintech).

### qPCR analysis

Total RNA was extracted using the Nucleospin RNA plus kit (Macherey Nagel, Hoerdt, France) and converted into cDNA using the Maxima First Strand cDNA synthesis Kit (Thermo Scientific, Illkirch, France). Quantitative RT-PCR (qPCR) was carried out using the EurobioGreen qPCR Mix Lo-Rox on a qTOWER instrument (Analytik-jena, Jena, Germany). Each reaction was conducted in a final volume of 10 μl containing 4 ng RNA equivalent of cDNA and 150 nM of primers. The primer sequences utilized for DNA amplification were for 5’-CCTCAGGAGTCCTTGTGCAG and rev 5’-GAACAGCAAGGTGGCCTTTG for *Oas3* and for 5’-GTGACAAGCCTGTAGCCCAC and rev 5’-GTACAACCCATCGGCTGGC for *Tnf*.

### Flow cytometry analysis

Cell death was assessed by staining the cells with Annexin V-FITC (from Miltenyi Biotec, Bergisch Gladbach, Germany) and propidium iodide (10 μg/ml) following the manufacturer’s instructions. Characterization of CD8+ T cells and NK cells was assessed by staining cells respectively with FITC-anti-murine CD8a (BD Biosciences 53-6.7; 1:100) and PE/Cya7-anti-murine NKp46 (BioLegend 29A1.4; 1:20). The flow cytometry analysis was conducted on a cytometer Accuri C6 plus (BD Biosciences). All experiments were conducted at least three times.

### Immunofluorescence

Cells were fixed with paraformaldehyde 4%, permeabilized with Triton 0.5% and stained with anti-p65 (Cell signaling 8242, 1:500) then with Alexa Fluor 488 anti-mouse (Thermo Fisher Scientific A11001, 1:500) as secondary antibody and with DAPI for nuclear immunolabelling (Thermo Fisher Scientific 62248, 1:2000). Mounted coverslips with cells were examined using 20X objective on the confocal AX LIPSI microscope (Nikon).

### 3’Sequencing RNA Profiling

Total RNA was obtained the Nucleospin RNA plus kit (Macherey Nagel, Hoerdt, France) according to the manufacturer’s instructions. RNA quality was verified with the Bioanalyzer System (Agilent Technologies) using the RNA Nano Chip and analyzed by 3’ Sequencing RNA Profiling (3’SRP). Digital gene expression profiles were generated by counting the number of UMIs associated with each RefSeq genes, for each sample. RNA-seq analysis were performed using the 3’SRP analysis pipeline developed by the BiRD core facility (https://doi.org/10.21203/rs.3.pex-1336/v1).

### In vivo experiments

Animal experiments were performed in accordance with protocols approved by the French regulations and the local ethics committee (APAFIS# 8629-2017011915305978 and # 36022-2022031016032320). The number of animals was estimated on the basis of experience gained with these models, taking into account tumor growth heterogeneity and post-operative complications, in order to obtain a sufficient number of mice at the end of the experiment for metastasis analysis. Orthotopic allografting consisted in injecting 40 μl of PBS containing 150,000 4T1-Luc2 cells into the fourth mammary fat pad of 7-week-old Balb/c mice. Primary tumors were surgically excised at D12 (early excision without metastatic spread) and D26 (late excision with metastatic spread) after allografting. Animals were sacrificed at D40. For the duration of the experiment, the animals were monitored by bioluminescence (intraperitoneal injection of 100 μl of D-Luciferin, 150 mg/kg, Interchim) to assess primary tumor growth before excision and to monitor recurrence and metastatic growth afterwards. The presence of endpoints was checked by observing rapid weight loss, weight loss greater than 20% of body weight, stooped posture, lethargy, absence of movement, rapid growth of tumor masses, mass greater than 1500 mm^3^, gait abnormalities, lesion interfering with eating and drinking, anuria or ulcerated tumor. Mice showing any of these signs were euthanized and excluded from analysis. Tumor diameters in length (L) and width (W) of primary tumors were measured every 2-3 days using a caliper. Tumor volume was calculated according to the following formula: (L^2*W)/2. Lung metastases were assessed at the end point by direct visual examination after euthanasia and by ex vivo bioluminescence to differentiate true lung metastases from pleural carcinomatosis. During in vivo experiments, mice were treated with: paclitaxel (Fresenius Kabi, Deutschland, 100 μl of product in intraperitoneal injection, 10 mg/kg once per week with 2 to 3 neoadjuvant and 1 adjuvant sessions) and/or navarixin (MedChemExpress, USA, feeding of 200 μl of product at 10 mg/kg 5 times per week with 3 weekly neoadjuvant and 1 adjuvant sequences). Control mice received the same amount of excipient without the therapeutic product (NaCl for Paclitaxel and corn oil for Navarixin).

### Tumor dissociation

Primary tumors were resected 12 days (early primary tumor, D12) or 26 days (late primary tumor, D26) after orthotopic allograft. Lung metastases were harvested 40 days after allograft. The various samples were immediately placed in DMEM medium on ice, then dissociated by mechanical (scalpel chopping) and enzymatic digestion (Collagenase A - from Clostridium histolyticum, Sigma-Aldrich, 1h at 37°C), within 2h of surgical resection or autopsy. After dissociation, cell suspensions were passed through a 70 μm filter, washed twice with medium and passed through erythrocyte lysis buffer (RBC Lysis Buffer 10X, Biolegend). The remaining cell suspensions were then resuspended in DMEM medium and enriched with viable cells using the Dead Cell Removal Kit magnetic sorting protocol (Miltenyi Biotec).

### ScRNAseq preparation

Single cell RNA sequencing was conducted using the Chromium Single-Cell 3’ v3.1 kit from 10X Genomics, according to the manufacturer’s protocol, multiplexing 2-4 mouse tumors or metastases per reaction according to the Chromium 3ʹ CellPlex Kit Set A protocol (PN-1000261). Libraries were sequenced on the NovaSeq 6000 platform (Paired-end, 28 bp Read1, 90pb Read2).

### Bioinformatic analysis

Raw BCL files were demultiplexing and mapped to the reference genome (mm10-3.0.0) using the Cell Ranger Software Suite (v.7.2.0). The filtered count matrices, containing only high quality barcodes, were loaded into Seurat objects using Read10X function (Seurat v.5.1.0) for further processing. Cells with fewer than 500 UMIs were discared as well as cells with more than 20 percent of mitochondrial genes. Doublets were identified using the scDblFinder R package (v.1.6.0), where the scDlbFinder score was calculated for each cell, and the scDblFinder threshold was applied for doublet identification. After preprocessing, normalization was performed using the NormalizeData function, with default parameters, which applies logarithmic normalization to gene expression data. Data were then centered and scaled using ScaleData function. During scaling step, regression of S and G2M phase scores was performed to minimized the influence of cell cycle effects in downstream analysis. The resulting data were projected onto an Uniform Manifold Approximation and Projection (UMAP) using the runUMAP function. Cell clustering was carried out with the FindNeighbors and FindClusters functions using varying resolutions to identify distinct clusters. For each defined cluster, gene markers analysis were performed using either the FindAllMarkers function which compares one cluster against all others, or the FindMarkers function which identifies differentially expressed genes between two conditions. Signature scoring using curated lists of genes was performed with the AddModuleScore function. Copy number variation analysis was performed using the InferCNV R package (v.1.16.0 https://github.com/broadinstitute/infercnv). The analysis was executed using the run function with the following parameters: denoise=TRUE, analysis_mode=”subcluster”,tumor_subcluster_partition_method=”leiden”leiden_resol ution=0.0015, , k_nn = 150. The normal reference set was defined using single-cell data from young adult mice as described by Carman Man-Chung et al (https://doi.org/10.1016/j.celrep.2020.108566). CellChat-v2 package (https://github.com/sqjin/CellChat) was used to investigate the interplay between tumor and immune cells. CellChatDB was used as reference network database. ComplexHeatmap (v.2.15.4), ggplot2 (v.3.4.4), dittooSeq (v.1.4.4) R packages were used for graphical representation.

### Transwell migration

Splenocytes collected from spleens of BALB/c mice were seeded in the top compartment of a Transwell chamber with 3-µm pore size (Corning, 3462). Three-hundred-micrometer-thick 4T1-Luc2 tumor sections (cut by vibrating blade microtome VT1200 S, Leica), freshly removed from Balb/c mice, were placed in the bottom compartment. After 48 h of incubation and treatment by paclitaxel and/or navarixin, cells in suspension from the bottom compartment were collected and numbers of viable and dead immune cells were calculated.

### Immunohistochemistry

Primary tumor and metastatic lung tissues were collected from Balb/c mice allografted with 4T1-Luc2 cell line. Three-micrometers-thick tissue sections of formalin-fixed, paraffin-embedded (FFPE) tumor tissues were treated with protease for 4min (Protease 1, Ventana, 760-2018) to achieve epitope retrieval. Samples were then incubated during 32min at 37°C with the following primary antibodies: Ki67 (BD Biosciences 550609, 1:500), CD8a (Abcam EPR21769; 1:2000), NCR1 (Abcam EPR23097-35; 1:100), Cleaved Caspase 3 (Cell Signaling 9131; 1:200). Protein - expression was detected with an OptiView DAB IHC Detection Kit (Roche Diagnostics, 760-700), optimized for automated immunohistochemistry (Benchmark GX stainer, Ventana Medical Systems, Roche Diagnostics). Ki67-immunolabelled tumor nuclei were quantified on 10 High Power Fields (HPFs, fields of 0.237 mm^2^ at 400X magnification). CD8a- and NCR1-immunolabelled cells were quantified on 10 HPFs in intratumor and peritumor (comprising a thickness of 200μm around the periphery of the tumor) areas. Cleaved caspase 3-immunolabelled areas were quantified with the ImageJ software on 10 HPFs and expressed in pixels per HPF.

### Statistical tests

GraphPad Prism was used to analyze data, draw graphs, and perform statistical analyses unless otherwise specified. Data were presented as means ± SEM. Statistical analyses were carried out by Student’s t test unless otherwise specified. *P ≤ 0.05, **P ≤ 0.01, and ***P ≤ 0.001 were considered significant. Representative results of three or more independent experiments were shown. Power analysis was conducted for all statistical studies. Blinding of sample names was used to reduce unconscious bias whenever possible.

## RESULTS

### Chemo-induced activation of STING-dependent IFN-I and STING-independent NF-κB secretory phenotype

We previously observed a major role in antitumor response to antimitotic chemotherapy (paclitaxel) of STING dependent paracriny using immunodeficient models of triple-negative breast cancer Lohard et al., 2020). We further explored the biological effects of paclitaxel stimulation on the secretory phenotype of TNBC cells using CRISPR-mediated STING knockout in 4T1-Luc2 murine cell lines (CRISPR^STING^ versus control cell line CRISPR^CTL^, Figure 1.A). Cytokine array on both cell lines revealed the necessity of constitutive STING expression for activation of type I interferon (IFN-I) pathway (as judged by the absence of CCL5 -a well-known interferon-stimulated gene (ISG)- secretion in CRISPR^STING^, Figure 1.B). *Oas3*, along with other ISGs, was transcriptionally reduced in CRISPR^STING^ cells, and was induced by in CRISPR^CTL^ cells (and not CRISPR^STING^ cells) exposed to paclitaxel, demonstrating STING-dependent chemo-induction of the IFN-I pathway (Figure 1.C). Strikingly, we observed that IκBα was consitutively phosphorylated to equal levels in both cell lines with similar nuclear expression of p65 protein by immunofluorescence analysis, indicating that baseline activation of canonical NF-κB pathway does not necessitate STING in these cells (Figures 1.D, 1.E). Moreover, we found that chemotherapy-induced overexpression of *Tnf* was unaffected by the absence of STING, indicating that paclitaxel can activate the NF-κB pathway regardless of STING expression (Figure 1.F). Tumor cell analysis revealed that the presence of STING enhanced chemo-induced cell death, since CRISPR^STING^ exhibited a significantly lower apoptotic rate following paclitaxel treatment compared to the control cell line (Figure 1.G), despites similar polyploidy rates (Suppl. Figure 1.A). Of note, the absence of STING was associated with increased expression of MCL1, an anti-apoptotic protein, prior treatment, which indicates that STING may help regulate the balance between cell death and survival by negatively modulating constitutive MCL1 expression (Figure 1.H). Treatment of both cell lines with the NF-κB inhibitor AS602868 (specific inhibitor of IKK2) tended to increase paclitaxel-induced cell death, suggesting that NF-κB activation contributes to tumor resistance to paclitaxel, independently of STING (Figure 1.I). NF-κB inhibition also led to a reduction in constitutive MCL1 expression (Suppl. Figure 1.B). Thus, STING enhances chemotherapy-induced cell death, while the NF-κB pathway, mechanistically unrelated to STING, contributes to resistance to paclitaxel while regulating anti-apoptotic mechanisms.

**Figure 1:**
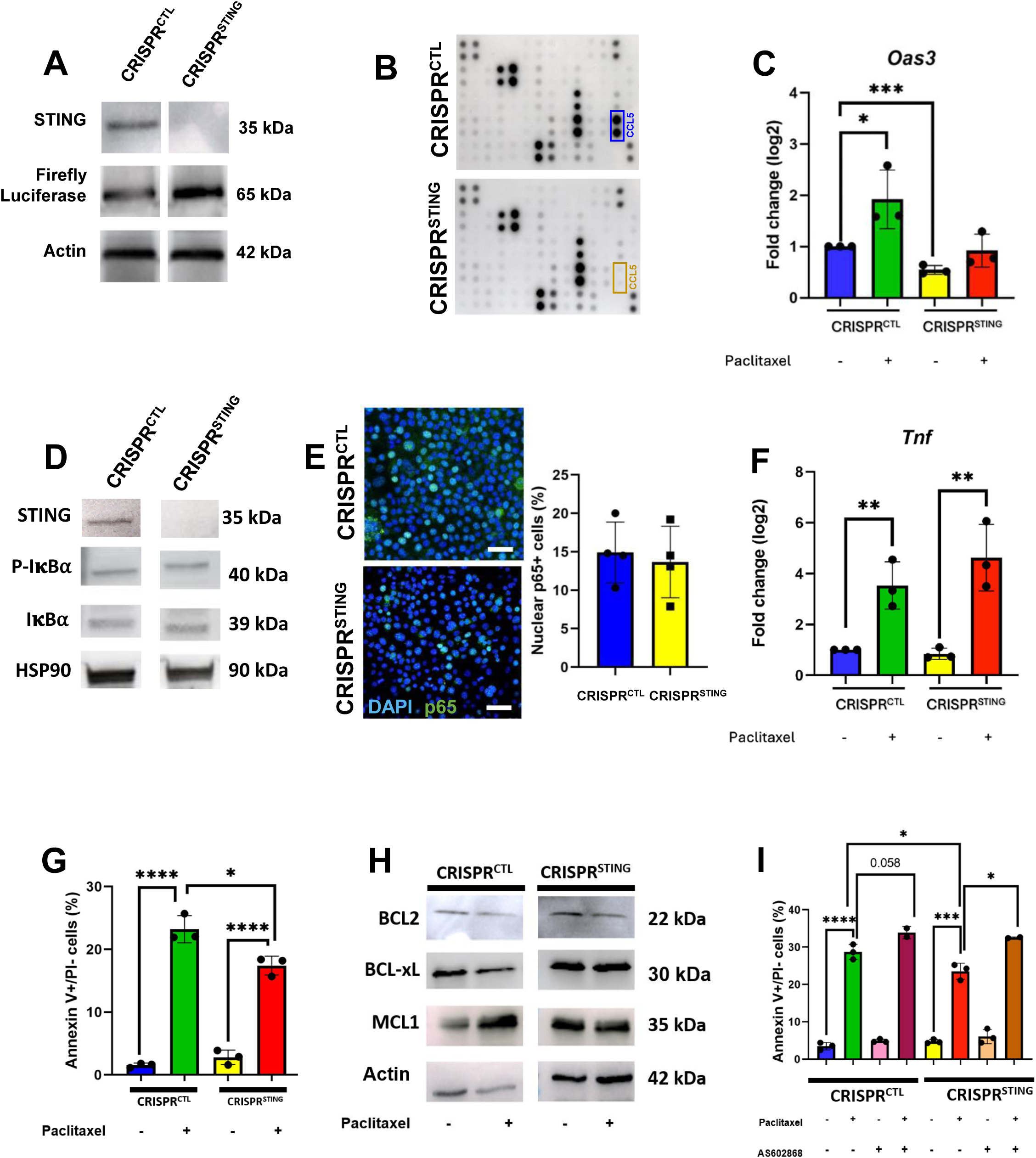
Chemotherapy induces both activation of an antitumor STING-dependent interferon pathway and a protumor STING-independent NF-kB pathway. **A:** Immunoblot analysis of STING and Firefly Luciferase expression in 4T1-Luc2 CRISPR^CTL^ and CRISPR^STING^. **B:** Cytokine array representing cytokines secreted in the culture medium from 4T1-Luc2 CRISPR^CTL^ and CRISPR^STING^. **C:** RT-qPCR analysis of *Oas3* in 4T1-Luc2 CRISPR^CTL^ and CRISPR^STING^ after paclitaxel treatment or not. **D:** Immunoblot analysis of the phosphorylation of IκBα in 4T1-Luc2 CRISPR^CTL^ and CRISPR^STING^. **E:** Percentage of nuclear p65+ cells by immunofluorescence in 4T1-Luc2 CRISPR^CTL^ and CRISPR^STING^; Scale bar: 50 micrometers. **F:** RT-qPCR analysis of *Tnf* in 4T1-Luc2 CRISPR^CTL^ and CRISPR^STING^ after paclitaxel treatment or not. **G:** Apoptotic cell death evaluated by flow cytometry of Annexin V+/Propidium Iodide- (AnnV+/PI-) cells in 4T1-Luc2 CRISPR^CTL^ and CRISPR^STING^ after paclitaxel treatment or not. **H:** Immunoblot analysis of anti-apoptotic proteins (BCL2, BCL-xL, MCL1) expression in 4T1-Luc2 CRISPR^CTL^ and CRISPR^STING^ after paclitaxel treatment or not. **I:** Apoptotic cell death evaluated by flow cytometry of AnnV+/PI- cells in 4T1-Luc2 CRISPR^CTL^ and CRISPR^STING^ after paclitaxel/AS602868 treatments or not.

We then sequenced the transcripts of both cell lines exposed or not to paclitaxel (3’Seq RNA Profiling) and identified upregulated and downregulated genes in each condition (Figure 2.A). This analysis confirmed that paclitaxel treatment significantly activated transcriptional signatures related to the NF-κB pathway in both cell lines regardless of STING presence (Figures 2.B, 2.C, 2.D, 2.E, Suppl. Figures 2.A, 2.B), predicting production of pro-inflammatory factors and chemokines (*Cxcl1, Cxcl5, Ccl20*…). This inflammatory phenotype is associated with upregulation of Epithelial-to-Mesenchymal Transition (EMT) and ER stress signatures and downregulation of apoptosis markers, suggesting a protumor phenotype favored by antimitotic chemotherapy. In contrast, STING pathway related to IFN-I signaling (*Ccl5*…) appeared less induced by paclitaxel. We next focused on CXCL1 and CCL5 secretion and found that paclitaxel-induced secretion of CCL5 was STING-dependent (Figure 2.F), while that of CXCL1 was not (Figure 2.G). Notably, CXCL1 secretion was highly dependent of NF-kB pathway, as evidenced by the loss of chemo-induced CXCL1 secretion after treatment with the inhibitor AS602868 (Figure 2.H). From a translational point of view, analysis of published clinical single cell expression data indicated that these two chemokines are expressed in subsets of human TNBC’s ecosystem (Suppl. Figure 2.C). These results confirmed the induction by chemotherapy of a mixed secretome from STING-dependent IFN-I pathway (CCL5) and from STING-independent NF-kB pathway (CXCL1).

**Figure 2:**
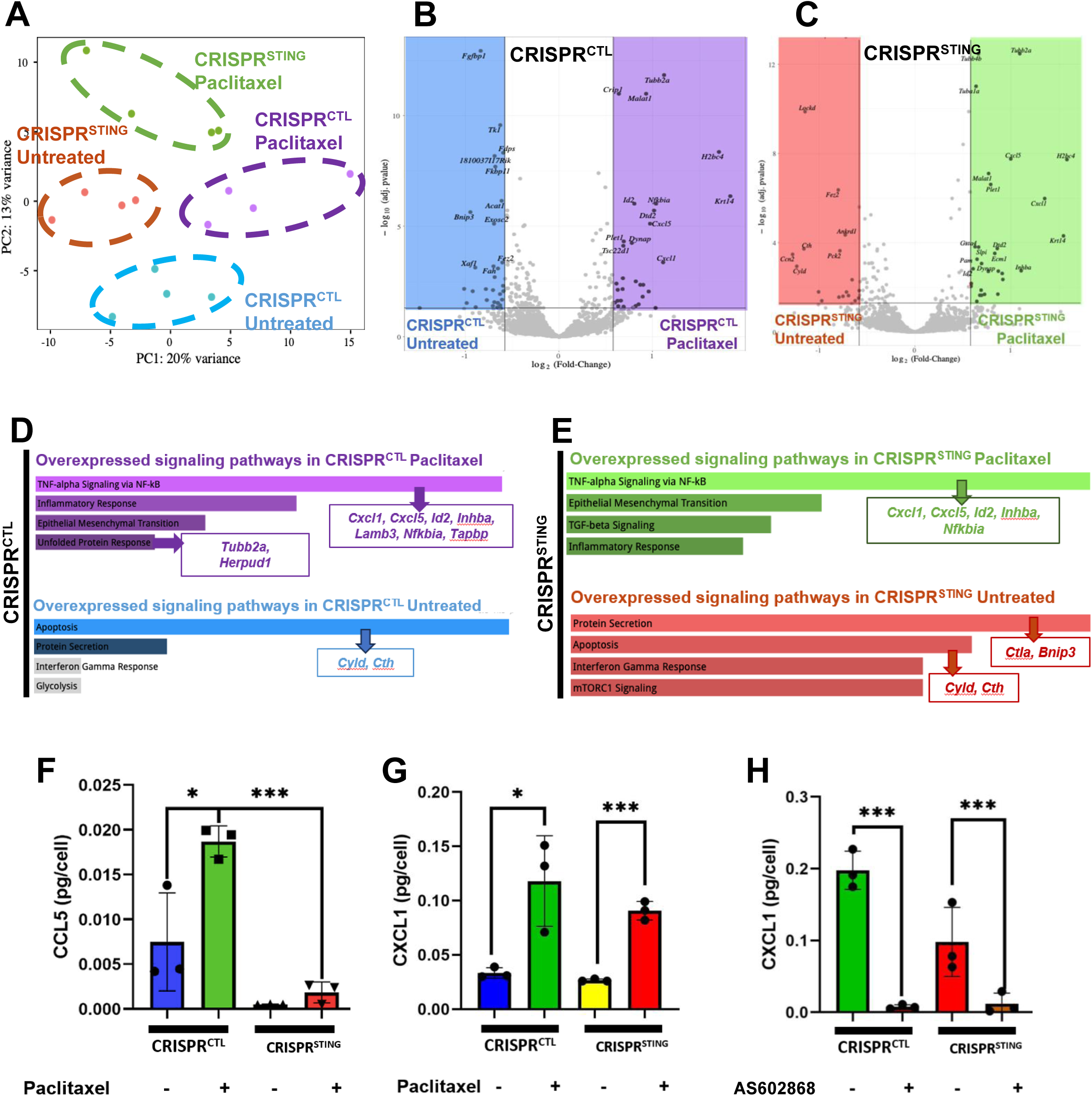
Chemo-induced activation of STING-dependent IFN-I and STING-independent NF-κB pathways are associated with tumor secretory phenotype. **A:** PCA analysis of 4T1-Luc2 CRISPR^CTL^ and CRISPR^STING^ treated or not by paclitaxel and analyzed by 3’Seq RNA Profiling. **B, C:** Volcano plots representing differentially up- and downregulated expressed under paclitaxel treatment in 4T1-Luc2 CRISPR^CTL^ and CRISPR^STING^, respectively. **D, E:** Signaling pathway signatures up- or downregulated by paclitaxel in 4T1-Luc2 CRISPR^CTL^ and CRISPR^STING^, respectively. **F, G:** Concentration in CCL5 and CXCL1 by ELISA in the culture medium of 4T1-Luc2 CRISPR^CTL^ and CRISPR^STING^ treated or not by paclitaxel. **H:** Concentration in CXCL1 by ELISA spot in the culture medium of paclitaxel-treated 4T1-Luc2 CRISPR^CTL^ and CRISPR^STING^ under treatment with AS602868 or not.

### Antimitotic chemotherapy alters the phenotype of tumor cells *in vivo*, favoring NF-kB signaling over IFN-I signaling during tumor progression

To study the impact of antimitotic chemotherapy on cancer cells within their ecosystem, during tumor progression, we set up a thorough experimental paradigm wherein we orthopically allografted 4T1-Luc2 cells in mammary fat pad of immunocompetent Balb/c mice and treated mice with neoadjuvant and adjuvant paclitaxel protocol at different stages of disease progression (Figure 3.A). We thus followed clinical evolution of mice and we analyzed tumor tissues at three different time points: early tumor without metastatic dissemination (d12), late tumor at a beginning metastatic stage (d26) and lung metastases (d40). Clinically, tumors appeared poorly sensitive to such paclitaxel treatment as treated mice showed no significant changes in tumor volume and metastases as untreated mice (Figures 3.B). In order to characterize the immediate impact of paclitaxel on tumor ecosystem, single-cell RNA sequencing (scRNAseq) was performed on tumor tissue (2-5 mice per condition). Tumor cells were isolated based on their positive expression of *Luc2* (coding for firefly luciferase, Suppl. Figure 3.A). The tumor origin was further confirmed by analyzing copy number variations (CNVs), which revealed strong CNVs in the *Luc2*^+^ cells. To better understand the phenotypic diversity within the tumor set, bioinformatic analyses were conducted to clusterize cells based on shared gene expression signatures on early tumor and applied on late tumor and metastases. Six main phenotypic clusters were identified (Label Transfer, Figures 3.C, 3.D, 3.E): (1) NF-κB cluster associated with genes constitutive of NF-κB signaling (e.g., *Nfkb1, Nfkbia*) and inflammatory chemokines (e.g., *Cxcl1, Cxcl2*); (2) proliferative cluster characterized by genes related to the cell cycle (e.g., *Birc5, Mki67*); (3) hypoxic cluster featuring genes involved in HIF- 1α induction and angiogenesis (e.g*., Vegfa, Mt1*); (4) basal cluster overexpressing basal keratins (e.g., *Krt7, Krt19*); (5) TGFβ cluster including genes involved in TGFβ signaling (e.g., *Smad3*) and (6) interferon cluster expressing interferon-stimulated genes (e.g., *Ccl5, Cxcl9, Ifit1*). Quantitative analysis of these clusters (Figure 3.F) confirmed that paclitaxel strongly selected for NF-κB signature cells, in the primary tumor and chronic exposure to chemotherapy in late tumors (3 neoadjuvant sessions of paclitaxel prior to late tumor removal) intensified this quantitative selection of NF-kB cluster. On the contrary, paclitaxel reduced all other phenotypic subgroups (hypoxic, basal and proliferative cells), the latter effect being consistent with its mitotic arrest properties and confirmed by decreased Ki67 index (Suppl. Figure 3.B). Thus, even if antimitotic chemotherapy had no significant clinical impact, it induced molecular changes in primary tumor populations by favoring cells with an inflammatory NF-κB signature. Of note, at the level of metastases where interferon cluster were absent, paclitaxel (1 post-removal adjuvant session) had no detectable effect on the proportion of different phenotypic clusters (Figure 3.F), possibly because our label transfer approach based on the tumor phenotypes of primary tumors blunted the detection of metastatic specific changes (see below).

**Figure 3:**
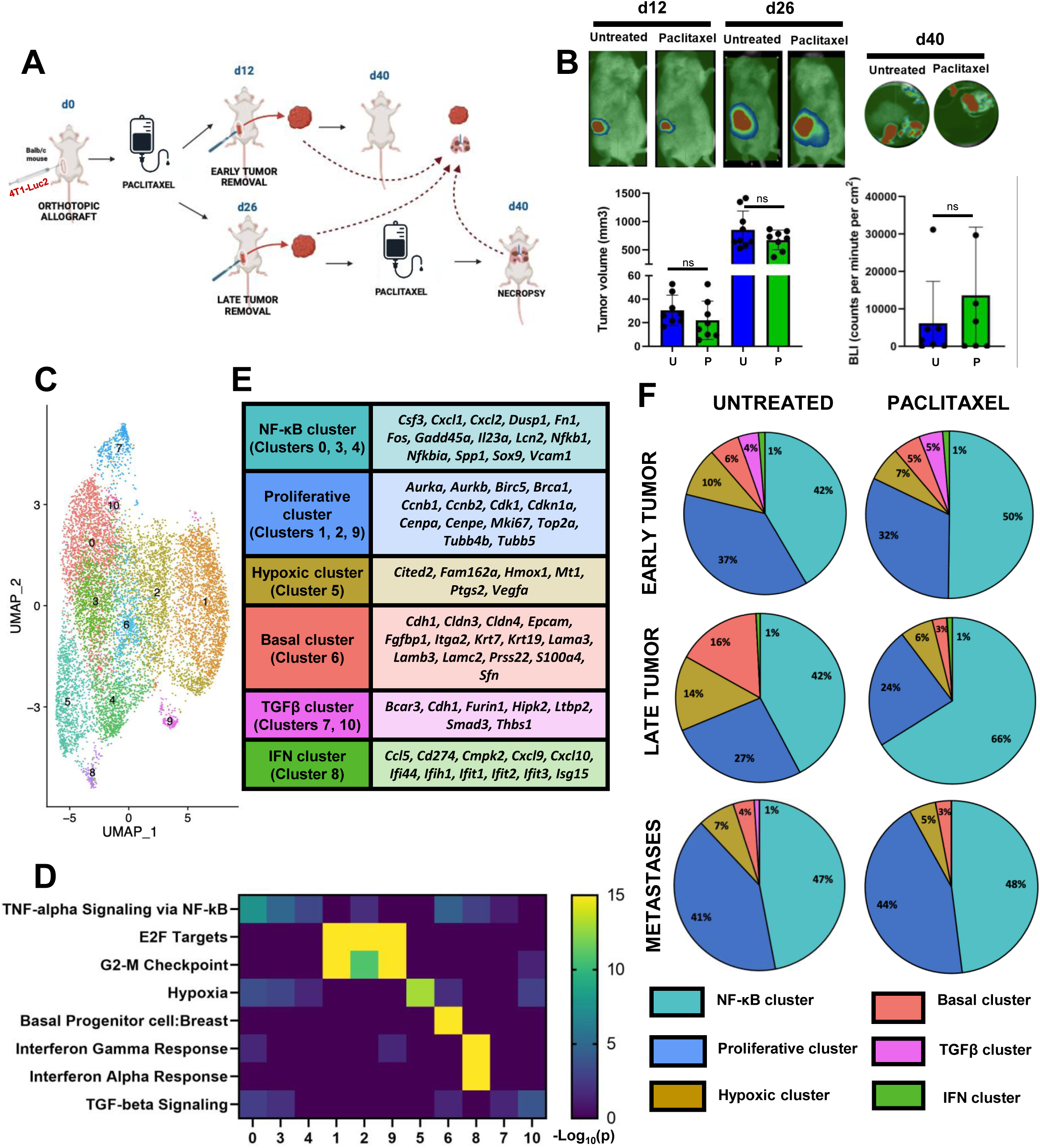
Chemotherapy alters the phenotype of tumor cells *in vivo*, favoring NF-kB signaling over IFN-I signaling. **A:** Experimental in vivo protocol mimicking tumor progression with orthotopic allograft (d0), paclitaxel neoadjuvant therapy (d7, d14, d21), early (d12) or late (d26) tumor removal, adjuvant paclitaxel administration (d33) and sacrifice (d40). **B:** Tumor volume measured on mice and illustrated by bioluminescence (d12 & d26) and lung metastases evaluated by bioluminescence intensity of *ex vivo* lungs (d40). **C:** UMAP representation of *Luc2*+ tumor cells in early tumors at resolution 0.4 highlighting 11 clusters with specific phenotypes. **D:** Characterization of the phenotype of tumor cluster expression with signature pathways. **E:** List of phenotype signatures with corresponding overexpressed genes. **F:** Proportion of different clusters in tumor tissues of early, late and metastatic 4T1-Luc2 tumors treated by paclitaxel or not.

### Tumor progression is associated with NF-κB secretory switch favoring cellular survival and metastasis

CNVs were inferred from scRNAseq data to characterize clonal selection pressures and further decipher how NF-κB activity might contribute (Figure 4.A). Analysis of these CNVs revealed four main clonal patterns (CP) with their own selection behavior (Figures 4.B, 4.C): (1) CP-A decreasing at all stages of tumor progression; (2) CP-B, restrained during tumor growth but not metastatic dissemination; (3) CP-C expanded during both tumor growth and metastasis and (4) CP-D disadvantaged in primary tumors but strongly increased in metastasis. Paclitaxel did not significantly alter the clonal composition of primary tumors (CP-A and CP-B) but tended to favor CP-C cells while reducing CP-D. Gene expression in distinct clonal patterns revealed a switch in secretory behavior (Figure 4.D). CP-A (expressing proliferative markers) and CP-B, both present early in the primary tumor expressed markers of both the IFN-I (*Cxcl9, Cxcl10, Ccl5*) and NF-κB_1_ (*Cxcl1, Csf3*) pathways, and gave way to CP-C and D expressing other markers of a new NF-κB_2_ (*Cxcl2, Il1b*) signaling pathway in metastases. These secretory changes were associated with enhanced expression of EMT markers and changes in anti-apoptotic genes (from *Bcl2* to *Bcl2l1* and *Mcl1*).

**Figure 4:**
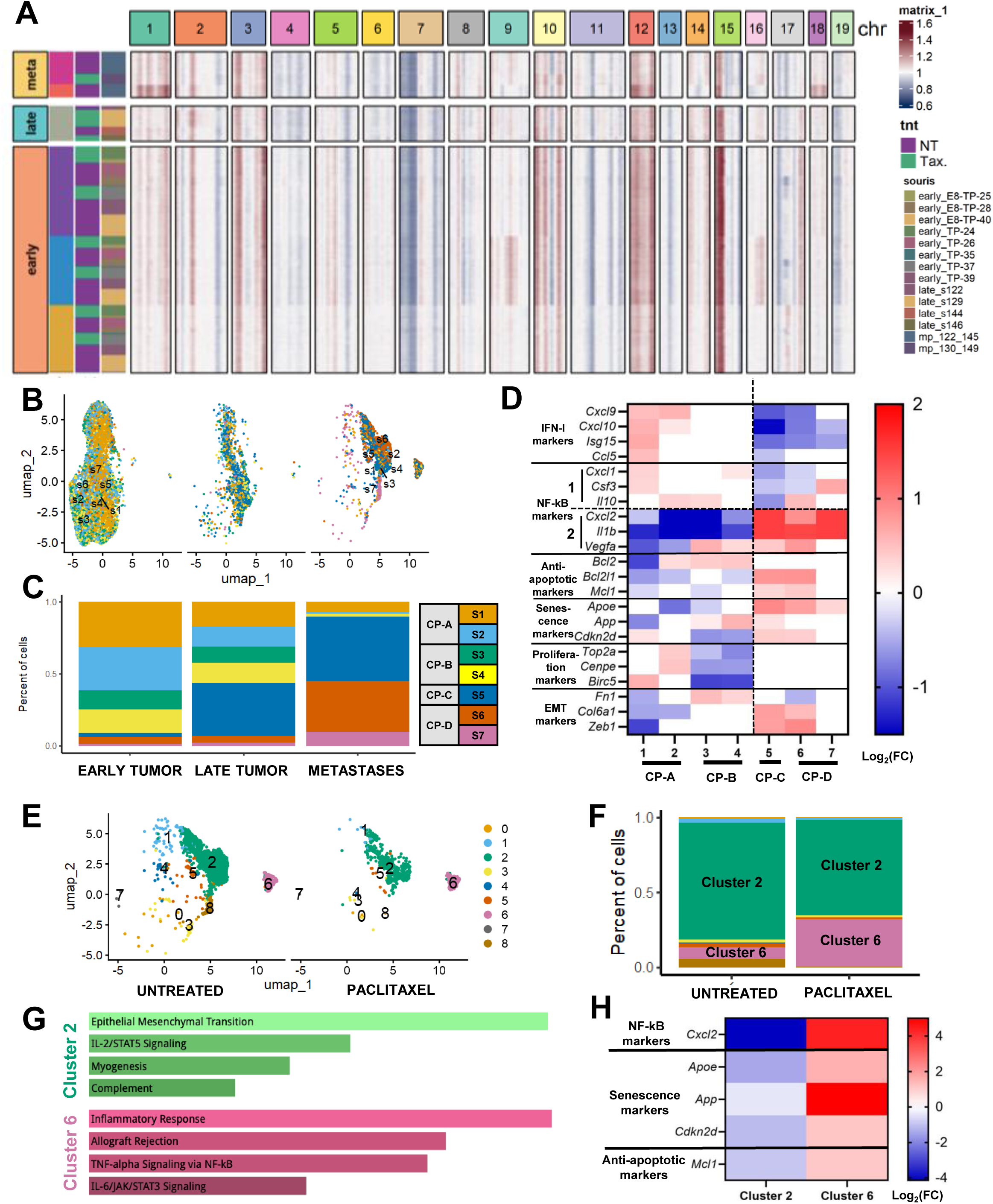
Tumor progression is associated with NF-κB secretory switch favoring cellular survival and metastasis. **A:** Heatmap showing inferred CNVs in cells of early, late and metastatic 4T1-Luc2 tumor tissues. **B:** UMAP representation of clonal clusters of 4T1-Luc2 cells in early, late and metastatic tumor tissues. **C:** Proportion of each clonal cluster and characterization of clonal patterns (CP) in early, late and metastatic tumor tissues. **D:** Heatmap showing gene markers of IFN-I, NF-κB, anti-apoptotic, senescence, proliferation and EMT signatures in each CP. **E:** UMAP representation of metastatic tumors treated by paclitaxel or not, characterized by phenotype clusterization. **F:** Proportion of each phenotype clusters in metastatic tissue treated or not by paclitaxel. **G:** Gene signature overexpression by the two mains phenotypic cluster in metastatic tissue (Clusters 2 and 6). **H:** Heatmap representing expression of Cxcl2, Apoe, App, Cdkn2d and Mcl1 in paclitaxel-treated metastatic clusters 2 and 6.

To characterize in more detail the impact of antimitotic chemotherapy on metastatic cells and better understand the mechanistic origin of their secretory switch, we isolated metastatic cells from our overall unsupervised clusterisation (Figures 4.E). We identified that two phenotypically distinct (that is showing distinct mRNA expression profiles: Cluster 2 and Cluster 6) accounted for the majority of metastases (Figure 4.E), with cluster 2 disfavored by paclitaxel, which benefited Cluster 6 (Figure 4.F). Cluster 2 overexpressed genes associated with an EMT signature, while Cluster 6 overexpressed genes related to the inflammatory response, particularly those linked to the NF-kB pathway (Figure 4.G). Importantly, these phenotypic differences are unlikely to ensue from clonal constrains as inferred CNV indicated similar clonal compositions in cluster 2 and 6 (Figure 4.B). Upon further analysis of the overexpressed genes, we found that Cluster 6 exhibited strong expression of the NF-κB_2_ secretory phenotype (*Cxcl2^high^*), which was associated with overexpression of *Mcl1* (Figure 4.H). This argues that a secretory response to paclitaxel in metastases, is provided by a subset of cells that have gained an inflammatory phenotype at the expense of a mesenchymal one.

### Tumor NF-kB phenotype is involved in neutrophil chemotaxis within the tumor ecosystem

The *Ptprc*^+^ (coding for pan-immune marker CD45) immune cells from dissociated murine tumor tissues, including early primary tumors and metastatic lung tissues, were analyzed separately from scRNAseq data (Suppl. Figure 4.A). Various clusters were identified within the immune compartments (Figures 5.A, 5.B, Suppl. Figure 4.B): five from lymphoid lineage (CD8+ T cells, CD4+ T cells comprising Tregs, NK cells, and B cells) and four from myeloid lineage (macrophages, conventional and plasmacytoid dendritic cells and neutrophils). The analysis revealed that in early tumors, lymphoid cells were dominant while myeloid cells predominated in metastatic microenvironment. Paclitaxel treatment favored the presence of CD4+ T cells and NK cells in the early tumors at the expense of CD8+ T cells, macrophages and neutrophils (Figure 5.C). In sharp contrast, metastases showed an increase proportion of both macrophages and neutrophils after paclitaxel treatment. The tumor NF-κB-driven tumor secretory phenotype was linked to the recruitment of neutrophils, specifically through overexpression of the *Cxcr2* receptor in our murine model as in TNBC patients (Suppl. Figures 4.C, 4.D). An analysis, using the CellChat tool, hinted on tumor NF-κB activity promoting neutrophil chemotaxis, and interferon cluster cells interacting with anti-tumor effector cells such as CD8+ T cells and NK cells (Figure 5.D), which could explain a link between the secretory switch observed during tumor progression and the increase in the myeloid/lymphoid cell ratio during tumor progression. To characterize the profile of chemo-attracted neutrophils in the metastatic microenvironment, we isolated and analyzed them at higher resolution that allowed identifying two neutrophilic sub-clusters, A and B, both quantitatively amplified by paclitaxel (Figure 5.E, Figure sup. 4.E). Neutrophil sub-cluster A showed a less secretory phenotype and lower expression of *Cxcr2* while sub-cluster B overexpressed *Cxcr2* and NF-kB chemokine genes, with paclitaxel enhancing this difference between clusters (Figure 5.F). Chemo-induced NF-κB-driven secretory phenotype along tumor progression was thus associated with myeloid cells concentration in tumor microenvironment, notably secreting *Cxcr2*^+^ neutrophils amplifying chemokine secretion.

**Figure 5:**
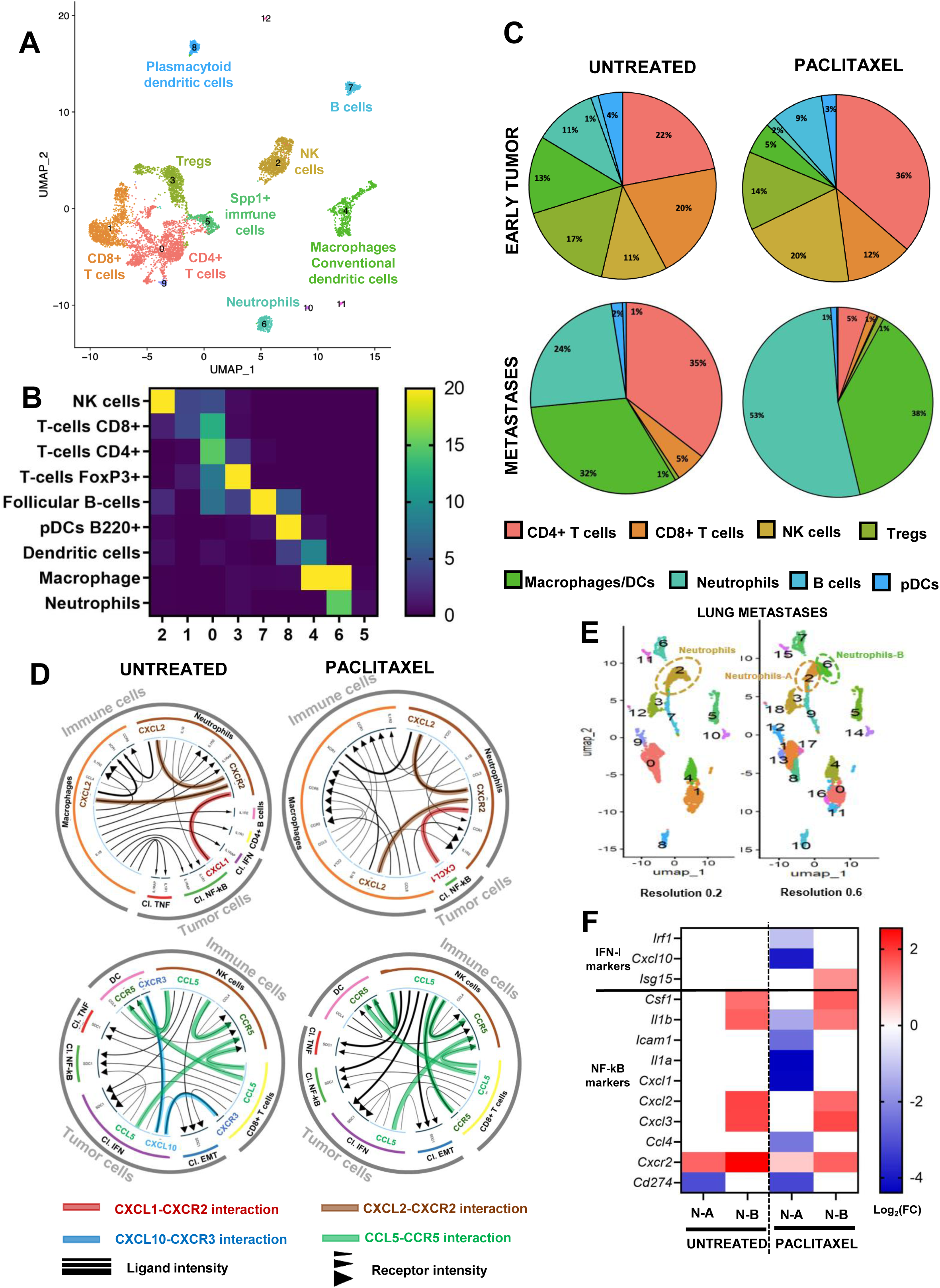
Tumor NF-kB phenotype is involved in to neutrophil chemotaxis within the tumor ecosystem. **A:** UMAP representation of *Ptprc*+ immune cells in early tumors at resolution 0.1 highlighting 9 mains clusters with specific phenotypes. **B:** Characterization of the phenotype of immune cluster expression with signature pathways. **C:** Proportion of different immune clusters in tumor tissues of early and metastatic 4T1-Luc2 tumors treated by paclitaxel or not. **D:** Cellular ligand-receptor cytokine interaction between tumor and immune cells of early tumors by CellChat analysis **E:** UMAP representation of metastatic immune microenvironment at resolution 0.1 and 0.6. **F:** Heatmap representing gene markers of IFN-I and NF-κB signatures by neutrophils-A (N-A) and neutrophils–B (N-B) in metastases under paclitaxel treatment or not.

### NF-κB-driven *Cxcr2*^+^ neutrophils are associated with an immunosuppressive microenvironment along tumor progression

In order to understand whether the decrease in effector cells in metastases was only the consequence of a decrease in the tumor interferon secretory phenotype, or whether NF-κB-driven chemo-attracted neutrophils also participated in this immunosuppressive microenvironment, we inferred interactions between neutrophils and the effector cells of antitumor immunity from scRNAseq data. Analysis of intercellular communications suggested that neutrophils, through the expression of PD-L1 (encoded by *Cd274*), may contribute to the inhibition of CD8+ T cell responses (Figure 6.A). Furthermore, the increase in neutrophil numbers during tumor progression was negatively correlated with NK cells numbers (Figure 6.B). Although chemotherapy increased the number of CD8+ T cells in the tumor microenvironment, particularly within the primary tumor itself, NK cells did not penetrate the tumor and remained mostly excluded in the periphery, as observed using Immunohistochemical analysis (Figures 6.C, 6.D). This immune exclusion was associated with an increase in exhausted cells (subgroups B) under paclitaxel treatment with no difference in the number of cytotoxic CD8+ T cells and NK cells (subgroups A overexpressing granzyme markers, (Figures 6.E 6F, 6G). All these results suggest an increasing immunosuppressive microenvironment associated with the presence of *Cxcr2*+ neutrophils.

**Figure 6:**
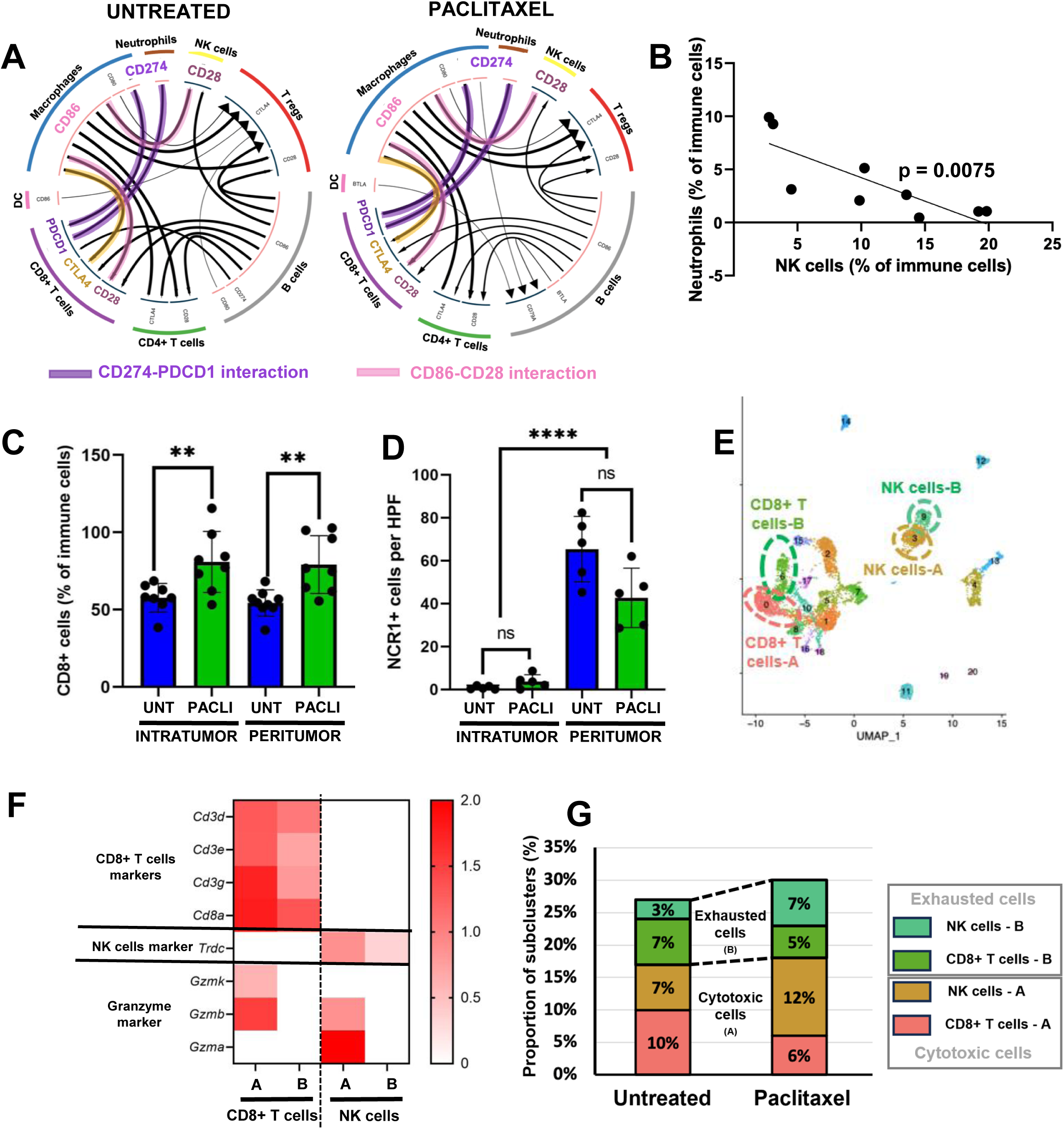
NF-κB-driven *Cxcr2*^+^ neutrophils are associated with an immunosuppressive microenvironment along tumor progression. **A:** Cellular ligand-receptor immune checkpoint interaction between immune cells of early tumors by CellChat analysis in paclitaxel or untreated cells. **B:** Proportion of Neutrophils and NK cells per mice in early tumors. **C, D:** Immunohistochemical count of intratumor and peritumor CD8+ T cells and NCR1+ NK cells in late tumors treated by paclitaxel or not. **E:** UMAP representation of immune microenvironment at resolution 0.8 in early tumors. **F:** Heatmap representing gene markers of CD8+ T cells and NK cells as well as granzyme genes in early tumors. **G:** Proportion of cytotoxic and exhausted CD8+ T cells and NK cells in early tumors.

### Inhibition of CXCL1/2-CXCR1/2 participates in reactivating antitumor response promoting tumor cell death

As TNBC cell subsets exhibited a chemo-induced NF-kB secretory phenotype contributing to an immunosuppressive tumor-promoting microenvironment, we explored the impact of pharmacological inhibition the CXCL1-2/CXCR1/2 pathway on both the immune response and tumor progression. We began by targeting CXCL1 using CRISPR/Cas9 to knock out the *Cxcl1* gene in the 4T1-Luc2 tumor cell line (Suppl. Figure 5.A). We found that knocking out CXCL1 did not affect tumor cell death or proliferation *in vitro*, suggesting minimal autocrine effects of this chemokine (Suppl. Figures 5.B, 5.C). In contrast *in vivo* modelization of CRISPR^CXCL1^ tumor-bearing mice showed reduced tumor volume compared to controls, confirming that CXCL1 plays a role in promoting tumor growth (Figure 7.A). As tumor cells express both CXCL1 and CXCL2 along the tumor progression, we investigated as an alternative strategy the effects of navarixin, a selective antagonist of both CXCR1 and CXCR2, which are receptors for several ELR+ CXC chemokines (Wu et al. 2022), including CXCL1 and CXCL2. Our analysis of human TNBC scRNAseq data revealed overexpression of these ELR+ CXC chemokines by TNBC cells, with two of them (*CXCL3*, *CXCL17*) associated with poor survival outcomes (Suppl. Figure 5.D). In our model, the combination of paclitaxel and navarixin significantly reduced tumor volume and metastatic spread, indicating the protumor impact of CXCL1/2-CXCR1/2 axis all along tumor progression (Figures 7.C, 7.D). To characterize the immune-related effects of navarixin, we analyzed immune cell migration in tumor tissues by placing tumor slices in culture under splenocytes to allow migration towards the tumor area after treatment with paclitaxel and/or navarixin (Figure 7.E). We found that paclitaxel had cytotoxic effects on immune cells, while navarixin did not, allowing viable immune cells to migrate into the tumor. Navarixin treatment attracted NK cells to the tumor site, in particular upon paclitaxel treatment, enhancing their infiltration into the tumor while CD8+ T cell migration was poorly affected. Immunohistochemical analysis of tumors confirmed that navarixin did not significantly affect CD8+ T cell migration but promoted intratumor NK cell infiltration, confirming an immunosuppressive impact of Cxcr2+ neutrophils on NK cells (Figure 7.F). The combination paclitaxel/navarixin also induced increased cell death in tumor cells, evidenced by cleaved caspase-3 immunolabelling on tumor slices (Figure 7.G). Thus, navarixin, by inhibiting the CXCL1-CXCR1/2 pathway and the chemotaxis of CXCR2+ neutrophils, reactivates the immune system, particularly through NK cell recruitment to the tumor site. When combined with paclitaxel, it enhances tumor cell death and reduces metastasis, providing a promising approach to modifying the tumor microenvironment and improving cancer treatment outcomes.

**Figure 7:**
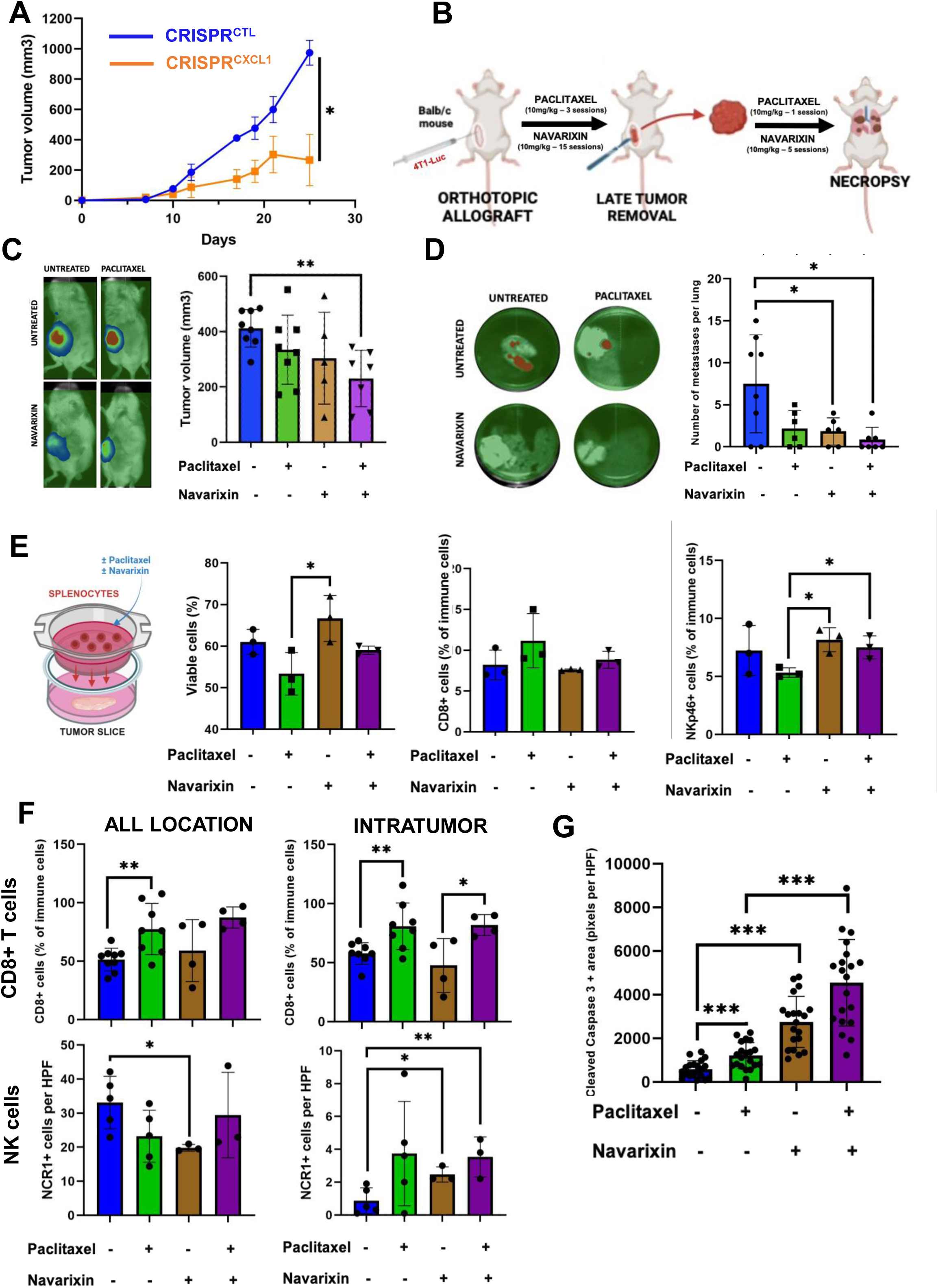
Inhibition of CXCL1/2-CXCR1/2 participates in reactivating antitumor response, and promotes tumor cell death. **A:** Primary tumor growth of 4T1-Luc2 CRISPR^CTL^ and CRISPR^CXCL1^ in Balb/c mice. **B:** Experimental *in vivo* protocol of Balb/c mice allografted with 4T1-Luc2 cell line and treated with paclitaxel and/or navarixin or not. **C:** Tumor volume measured on mice treated with paclitaxel and/or navarixin or not (d26). **D:** Number of lung metastases on mice treated with paclitaxel and/or navarixin or not (d40). **E:** Characterization (viability, CD4 and NKp46 markers) of splenocytes migrating across a transwell membrane in the culture medium containing ex vivo 4T1-Luc2 tumor slices and treated with paclitaxel and/or navarixin or not. **F:** Immunohistochemical count of CD8+ T cells and NCR1+ NK cells (all location and intratumor) in tumors treated by paclitaxel and/or navarixin or not. **G:** Immunohistochemical immunolabelling of cleaved caspase 3 in 4T1-Luc2 tumors treated or not by paclitaxel/navarixin.

## DISCUSSION

We previously demonstrated that antimitotic chemotherapy induces micronuclei accumulation in cancer cells, promoting the activation of the cGAS/STING pathway. Most relevantly, this pathway triggered a secretome linked to type I interferon and TNF pathways, inducing the NOXA expression in neighboring cells, making them particularly sensitive to BCL-xL inhibition (notably by BH3-mimetics, Lohard et al. 2020). These results highlighted a paracrine secretory role of the cGAS/STING pathway, promoting antitumor response by enhancing apoptotic cell death in an immunodeficient context. Other studies have also shown the crucial role of the secretome induced by STING in the antitumor response, in connection with the immune microenvironment, through chemotaxis and activation of antigen-presenting cells and effector immune cells (cytotoxic T cells and NK cells, An et al., 2019; Gan et al., 2022; Holicek et al., 2024). The idea of administering STING agonists as immunotherapy in combination with chemotherapy has thus gained attraction in the scientific community, aiming to potentiate the immunogenic death of tumor cells (Wang et al., 2022; Zhu et al., 2024). However, clinical trials have shown only a modest survival benefit for these therapeutic protocols (Meric-Bernstam et al., 2022). Recent studies suggest that the chronic activation of cGAS/STING pathway may also contribute to tumor progression by promoting immunosuppression, exhaustion of effector immune cells and metastatic spread (Bakhoum et al., 2018; Hong et al., 2022; Li et al., 2023; Zhao et al., 2024; Filderman et al. 2024).

Some authors proved that this poor prognosis induced by cGAS/STING pathway in TNBCs was due to NF-kB pathway activation and release of pro-inflammatory cytokines such as IL-6 (Hong et al., 2022). Our data do not dispute this, and cannot exclude that STING and NF-κB pathways cooperate *in vivo* to promote tumor progression. However, our results unambiguously establish that chemo-induced NF-kB pathway does not need STING expression to be patent. Possible pharmacological modulation of this protumor pathway could therefore help to reduce cancer progression (Poma et al. 2017). However, the NF-kB pathway is essential to many physiological cellular mechanisms (immunity, skin development, metabolism…), and these therapies have so far been associated with numerous side effects (Ramadass et al., 2020).

Some authors recently explored inhibition of alternative chemo-induced mechanistic stimulation of NF-kB pathway, notably the neutralization of IL-1a cytokine released from paclitaxel-induced dead cells diminishing IL-1R-MyD88-dependent NF-kB secretory phenotype (Hänggi et al. 2024)

We showed that tumor progression associated with repeated administration of antimitotic chemotherapy leads to a decrease in the type I interferon phenotype in favor of an NF-kB-dependent secretory phenotype. This phenomenon, described as tachyphylaxis, was demonstrated *in vitro* by a reduction in the secretion of ISGs (*OAS3, CXCL10, CCL5, ISG15*) after repeated STING pathway stimulations (poly:IC), while these same stimulations contributed to an increase in markers of STING-dependent NF-kB pathway, including *TNF* (Li et al., 2023). These authors attributed this increase to ER stress mechanisms, particularly EIF2α (*HSPA5*) and XBP1 pathways, while our results showed that, although there was a slight overexpression of *Hspa5* in our murine tumor cells with an NF-κB profile under paclitaxel, this ER stress stimulation was not observed permanently in the metastases. At metastatic sites, NF-κB activity seemed to be restrained to an overtly inflammatory subpopulation, distinct from a population endowed with mesenchymal traits. While the presence of the latter is in line with the notion of a role of EMT in metastasis, the presence of the former, which is favored by chemotherapy is less expected. We speculate that metastatic cells may transit back and forth between a “Mesenchymal” and an “Inflammatory” state. The molecular basis of this Mesenchymal to Inflammatory Transition (MIT) remains to be determined but it might play a major role in metastatic response to treatment given its immunosuppressive effects.

Recent publications have shown that senescent tumor cells, which initially suppress tumor growth, if not quickly cleared by the tumor immune response, can promote the proliferation, migration, and metastasis of surrounding cells (Guccini et al., 2021). Several observations suggest that cellular senescence could contribute to chemo-induced NF-κB activity: (1) NF-κB active clusters express markers of senescence, including known actors of a senescence-associated secretory phenotype (SASP, Figure 4.H); (2) *in vitro* paclitaxel treatment increase CXCL1-producing SA-β-galactosidase + cells (Suppl. Figure 6.A) and (3) published meta-analysis data show that CXCL1/2 chemokine expression, highly expressed in TNBCs correlates with the expression senescence markers (Suppl. Figures 6.B, 6.C, 6.D). NF-κB active cells, and their immunosuppressive effects, may thus be counteracted by senolytic agents, such as BH3 mimetics targeting BCL-2 family members (Malayaperumal et al. 2023; Zhu et al., 2016). In our model as in meta-analysis of human TNBCs, tumor progression was associated with overexpression of both senescence and pro-survival markers, notably the *Mcl1* transcript in metastasis. One recent publication highlighted the impact of MCL1 protein in the resistance to taxane therapy of senescent prostate cancer cells with increased apoptosis and reduced SASP after S63845 treatment (BH3-mimetics targeting MCL1, Troiani et al. 2022). However, administration of S63845 did not potentiate *in vitro* chemo-induced death of our murine tumor cells, probably due to the low efficacy of this molecule in targeting the murine form of MCL1 (Kotschy et al. 2016).

We also showed that secretory phenotypes changed upon chemotherapy-treatment during tumor progression, favoring *Cxcl1* expression in the primary tumor versus *Cxcl2* in metastases. Both secretory phenotypes participate in the attraction of neutrophils and the angiogenesis, and are known to promote tumor progression (Han et al., 2022; Zhang et al., 2018). However, CXCL1 would be more involved in inducing local immunosuppression enabling tumor growth (Bianchi et al., 2023; Wang et al., 2022), whereas CXCL2 would be more involved in remodeling the extracellular matrix and preparing the pre-metastatic niche through dialogue with tumor-associated macrophages and cancer-associated fibroblasts (Bao et al., 2022; Nambiar et al., 2023).

We further highlighted that tumor secretory phenotype was strongly involved in neutrophil chemotaxis within the tumor microenvironment, in particular neutrophils expressing the CXCR2 receptor. Numerous studies have focused on the protumor role of these cells favoring tumor growth, migration and participating to metastatic niche and disseminated cell implantation (Cheng et al., 2021; Lazennec et al., 2024). In particular, authors have characterized the different types of neutrophils, with antitumor IFN-stimulated neutrophils decreasing in cancer models (Zilionis et al., 2019; Benguigui et al., 2024). As in our model, this decline was in favor of neutrophils secreting pro-inflammatory cytokines (Masucci et al., 2019), but overexpression of the PD-L1 checkpoint protein (Wang et al., 2017) or release of metalloproteinases into the connective stroma (Sheng et al., 2023; Han et al., 2024), have also been suggested to promote tumor progression and resistance to treatment. We did not analyze the impact of neutrophil extracellular traps in our study, but authors have shown that neutrophils, by this way, promote progression as well as resistance to chemotherapy (Teijeira et al., 2020; Mousset et al., 2023).

Finally, we have shown that inhibition of the CXCL1/2-CXCR1/2 axis helped slow cancer progression in both primary tumors and metastases. TNBC is the breast cancer with the highest expression of CXCR2 within its immune microenvironment (Boissière-Michot et al., 2020), but the prognostic value of this marker remains controversial, with some studies associating CXCR2 with greater tumor infiltration and a better prognosis (Boissière-Michot et al., 2021), and others showing that it promotes stemness and favors tumor progression (Zhang et al., 2023). In our analyses, we have seen that different neutrophil types (both highly and mildly inflammatory) both overexpressed *Cxcr2*, suggesting a possible dual role for *Cxcr2*+ neutrophils. We note that navarixin is a dual inhibitor of both CXCR1 and CXCR2 receptors but *Cxcr1* was not highlighted in our scRNAseq analysis. This treatment has recently been the subject of a phase II randomized clinical trial involving patients with colorectal, lung and prostate cancers, in combination with pembrolizumab, but although associated with very few side effects, did not show any major efficacy in terms of survival (Armstrong et al., 2024). This trial did not include chemotherapy treatment, which we have shown to promote the expression of ligands promoting CXCR2+ neutrophil chemotaxis, and did not involve patients with TNBC.

In summary, this study highlights the major impact of STING-independent NF-κB pathway in the response of triple-negative breast cancer (TNBC) to antimitotic chemotherapy. We observed that, while the activation of the STING-dependent IFN-I pathway enhances chemotherapy-induced cell death, its effect decreases as cancer progresses; in contrast, the NF-κB pathway, which operates independently of STING, increases and contributes to tumor resistance by promoting anti-apoptotic mechanisms and creating a pro-inflammatory microenvironment. This secretory phenotype attracts CXCR2+ neutrophils, linked to an immunosuppressive environment that promoted tumor progression and metastasis. The combination of paclitaxel and navarixin significantly reduced tumor growth and metastasis, suggesting that targeting the CXCL1/2-CXCR1/2 pathway could improve therapeutic outcomes in TNBC by modulating the tumor immune landscape. Overall, these findings provide valuable insights into the molecular mechanisms underlying chemotherapy resistance and highlight a potential strategy to enhance the efficacy of cancer treatments.

## ACKNOWLEDGEMENTS

We thank Floriane Briand and Quentin Batard for their help during *in vivo* experiments, Julie Roul and Arulraj Nadaradjane for his assistance during different steps of scRNAseq protocol and Aurélie Fétiveau for her technical support to produce KO contructs. We thank Frédérique Nguyen for reviewing the article. We thank BioCore (UMS16 Inserm-UAR 3556 CNRS) biotechnology center, including Cytocell, Geno-A, PES (UTE-IRS-UN) core facilities, for technical support.

## AUTHORS CONTRIBUTION

F.C., V.J. and S.B.N. conducted experiments. F.C. and S.B.N. designed the experiments. F.C., F.G., P.P.J. and SBN. analyzed the data. E.D. gave technical assistance. F.C., P.P.J. and S.B.N. wrote the paper. P.P.J. and S.B.N. obtained fundings. P.P.J. and S.B.N. conceived the study and supervised it

## LEGENDS TO SUPPLEMENTARY FIGURES

**Supplementary Figure 1:**
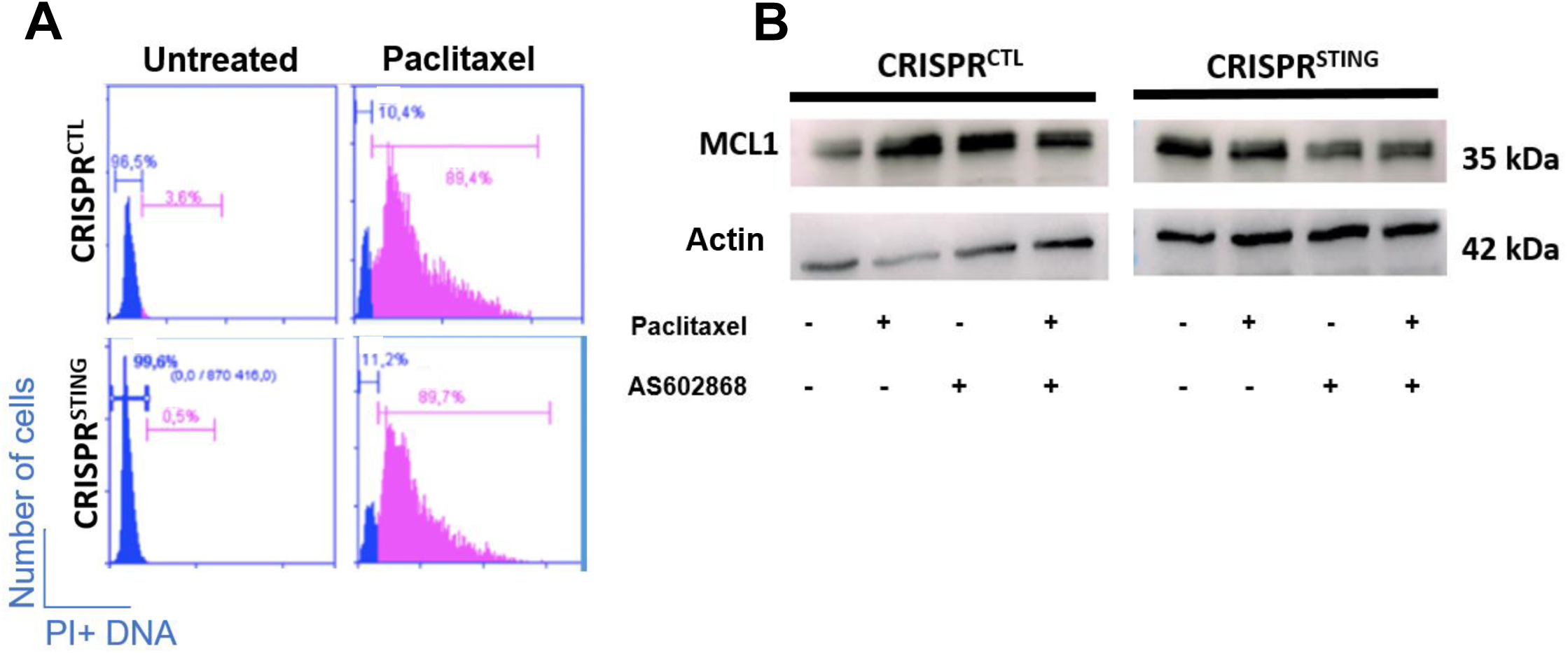
Characterization of DNA content and anti-apoptotic protein MCL1 of 4T1-Luc2 CRISPR^CTL^ and CRISPR^STING^ under paclitaxel treatment. **A:** Characterization of ploidy using PI DNA staining by flow cytometry in 4T1-Luc2 CRISPR^CTL^ and CRISPR^STING^ treated by paclitaxel or not. B: Immunoblot analysis of MCL1 expression in 4T1-Luc2 CRISPR^CTL^ and CRISPR^STING^ after paclitaxel and/or AS602868 treatments or not.

**Supplementary Figure 2:**
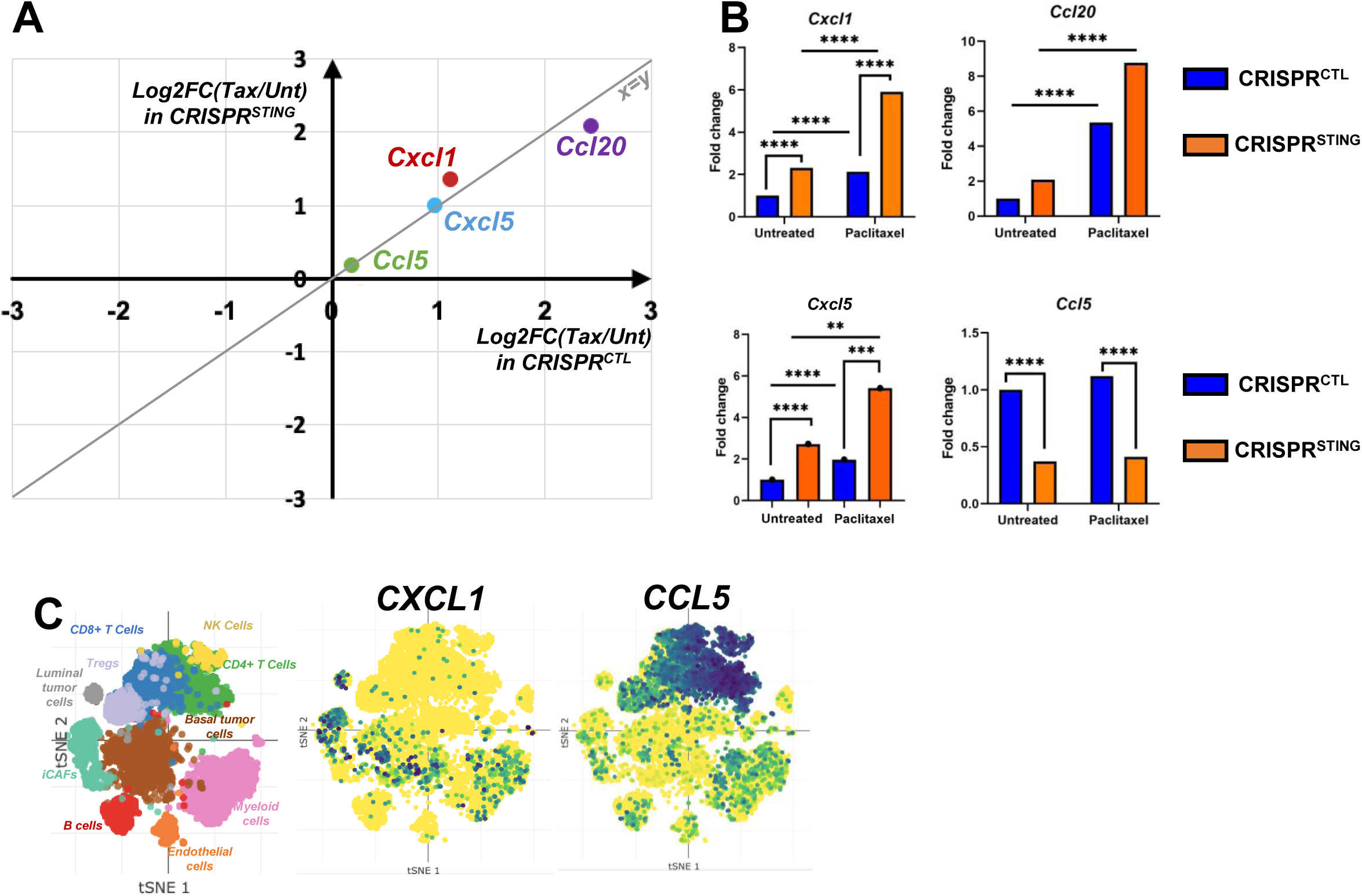
Characterization of secretory phenotype of 4T1-Luc2 CRISPR^CTL^ and CRISPR^STING^ under paclitaxel treatment. **A:** Impact of STING on paclitaxel-induced transcriptional expression (3’SRP) of *Cxcl1, Cxcl5, Ccl5* and *Ccl20* comparing fold-change expression (paclitaxel versus untreated) in both CRISPR^CTL^ and CRISPR^STING^. **B:** Transcriptional expression fold change of *Cxcl1, Cxcl5, Ccl5* and *Ccl20* in CRISPR^CTL^ and CRISPR^STING^ under paclitaxel or not. **C**: Expression of *CXCL1* and *CCL5* by human TNBC cells using scRNAseq analysis from Broad Institute (https://singlecell.broadinstitute.org/single_cell/study/SCP1106/stromal-cell-diversity-associated-with-immune-evasion-in-human-triple-negative-breast-cancer).

**Supplementary Figure 3:**
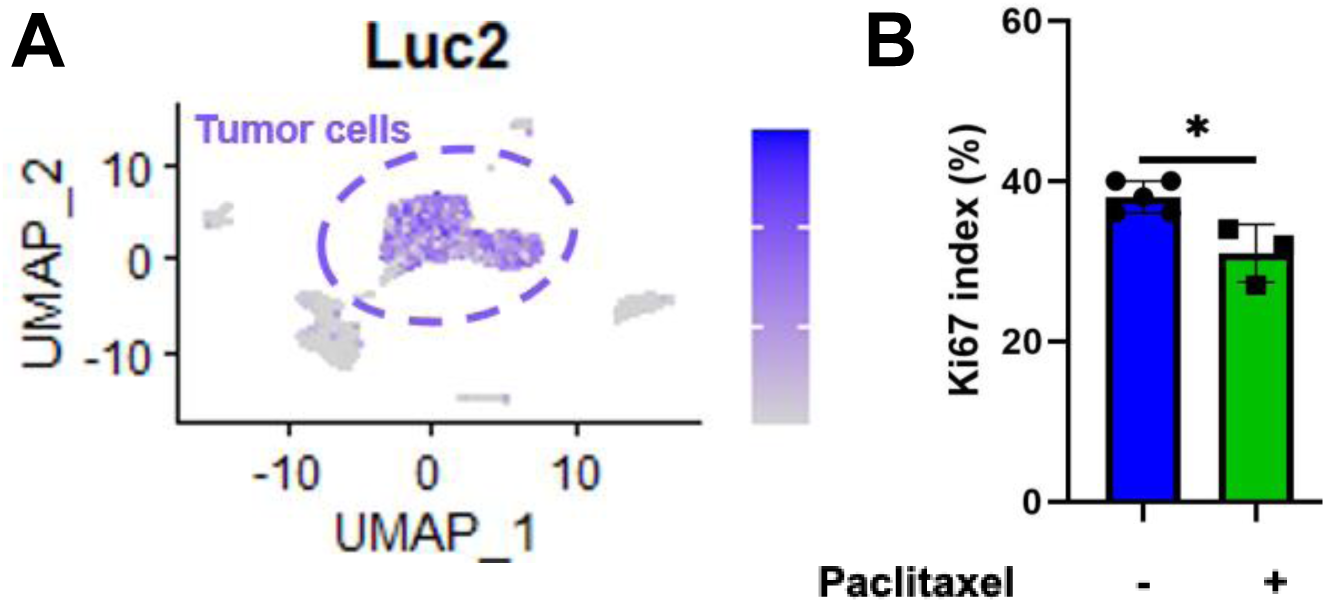
Characterization of tumor cells from in vivo experiments by scRNAseq analysis. **A:** Expression of *Luc2* marker by tumor cells in sequenced early tumors. B: Immunohistochemical Ki67 index of early 4T1-Luc2 tumors.

**Supplementary Figure 4:**
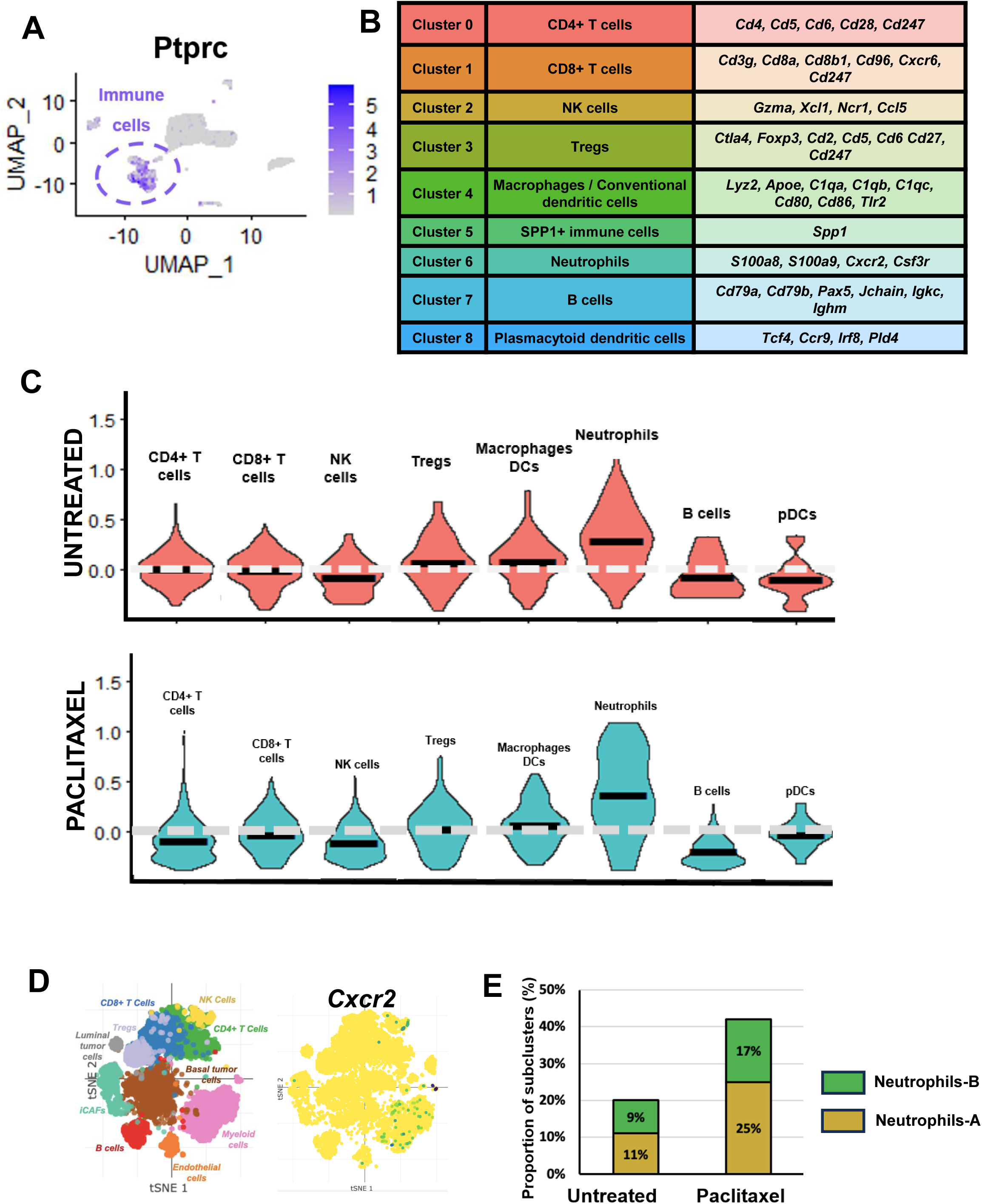
Characterization immune cells of 4T1-Luc2 microenvironment under paclitaxel treatment in scRNAseq. **A:** Expression of *Ptprc* marker by immune cells cells in sequenced early tumors. **B:** List of genes characterizing each immune cluster at resolution 0.1 in early tumors. **C**: Expression receptor signature specific of NF-κB secretory phenotype in immune cells of early tumors treated by paclitaxel or not. **D**: Expression of *CXCR2* by human TNBC cells using scRNAseq analysis from Broad Institute. **E**: Proportion of neutrophils-A and –B in metastatic 4T1-Luc2 tissue after paclitaxel treatment or not.

**Supplementary Figure 5:**
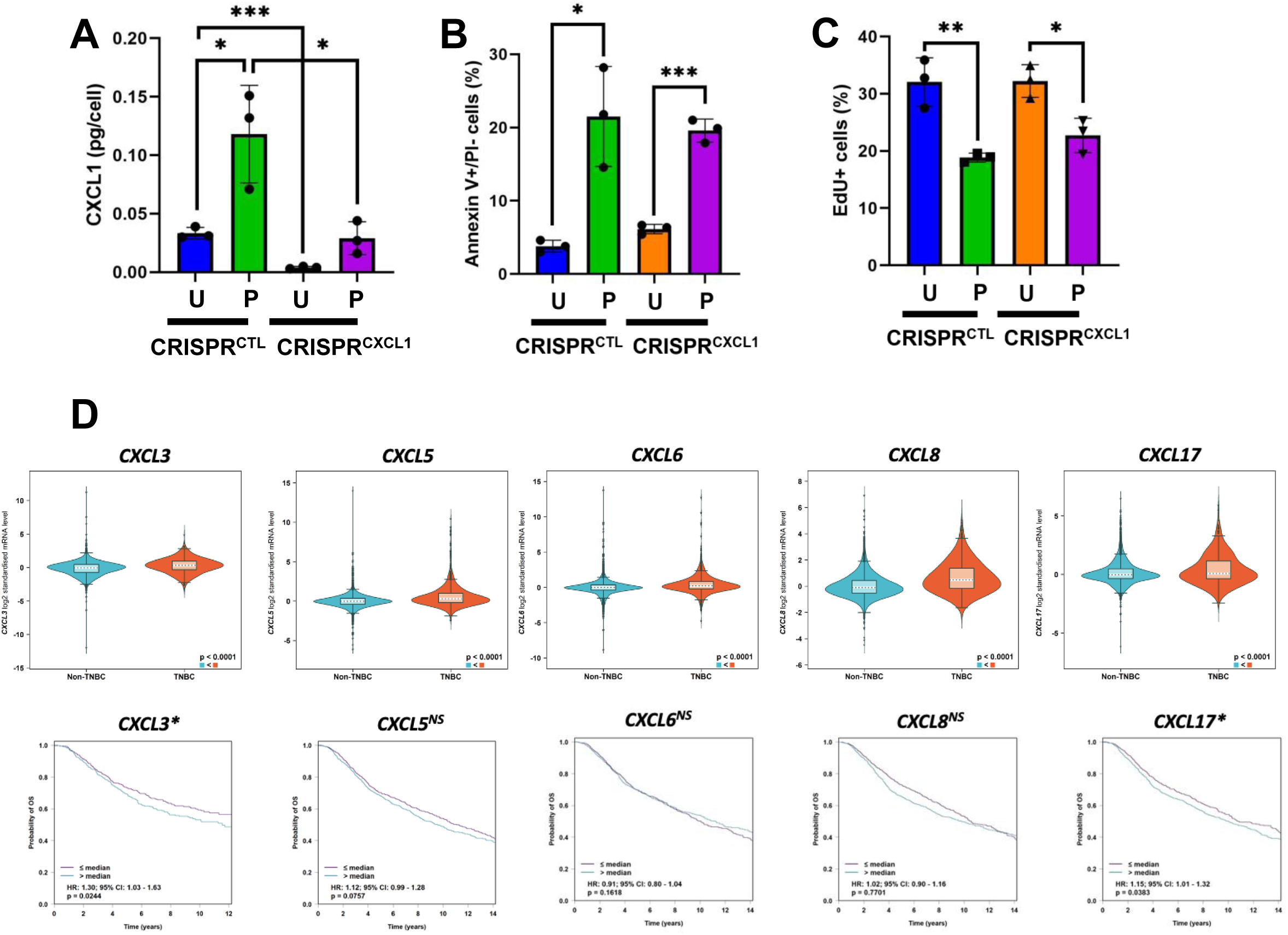
Characterization of the impact CXC ELR+ ligands on murine and human TNBCs. **A:** Concentration in CXCL1 of culture medium containing 4T1-Luc2 CRISPR^CTL^ and CRISPR^CXCL1^ cells treated by paclitaxel or not. **B:** Apoptotic cell death evaluated by flow cytometry of AnnV+/PI- cells in 4T1-Luc2 CRISPR^CTL^ and CRISPR^CXCL1^ after paclitaxel treatment or not. **C**: Proliferative EdU+ cells evaluated by flow cytometry of AnnV+/PI- cells in 4T1-Luc2 CRISPR^CTL^ and CRISPR^CXCL1^ after paclitaxel treatment or not. **D**: Expression of *CXCL3, CXCL5, CXCL6, CXCL8* and *CXCL17* (CXC ELR+ chemokines) by human TNBC cells and associated prognosis using BC-GenExMiner tool (Jézéquel et al., 2012; Jézéquel et al., 2013; Jézéquel et al., 2021).

**Supplementary figure 6:**
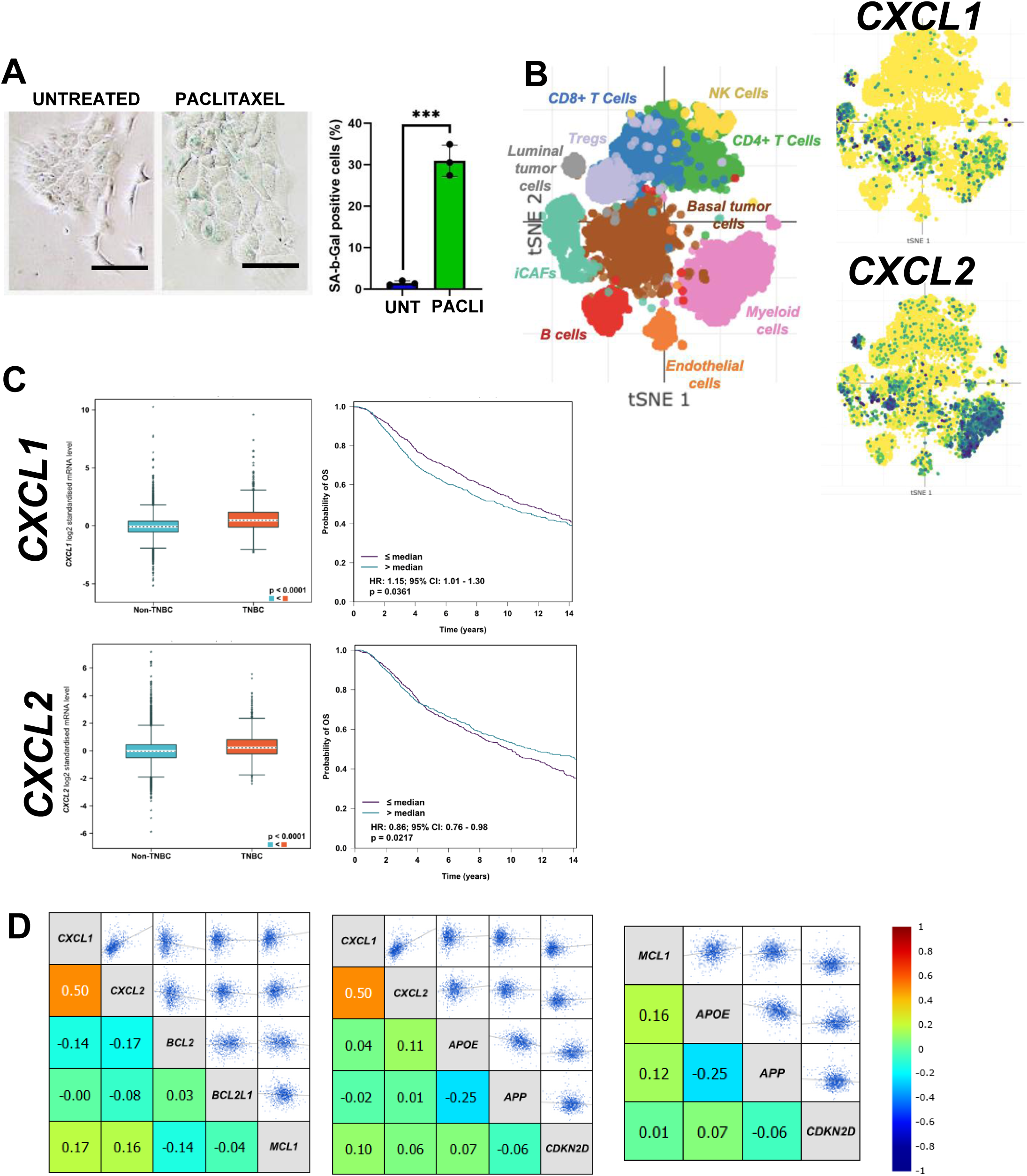
Senescence associated phenotype of 4T1-Luc2 under paclitaxel treatment in scRNAseq. **A:** SA-β-Gal reaction of 4T1-Luc2 cells treated by paclitaxel or not. **B**: Expression of *CXCL1* and *CXCL2* by human TNBC cells and their immune microenvironment (from scRNAseq of Broad Institute) **C:** *CXCL1* and *CXCL2* expression by human TNBC and associated prognosis according to the Bc-GenExMiner tool. **D**: Correlation between *CXCL1*/*CXCL2,* anti-apoptotic and senescence markers using BC-GenExMiner tool.

